# Nuclear stability and transcriptional directionality separate functionally distinct RNA species

**DOI:** 10.1101/005447

**Authors:** Robin Andersson, Peter Refsing Andersen, Eivind Valen, Leighton J. Core, Jette Bornholdt, Mette Boyd, Torben Heick Jensen, Albin Sandelin

**Affiliations:** The Bioinformatics Centre, Department of Biology and Biotech Research and Innovation Centre (BRIC), University of Copenhagen, Ole Maaloes Vej 5, DK-2200 Copenhagen, Denmark.; Centre for mRNP Biogenesis and Metabolism, Department of Molecular Biology and Genetics, C.F. Møllers Alle 3, Bldg. 1130, DK-8000 Aarhus, Denmark; Department of Informatics, University of Bergen, Thormøhlensgate 55, N-5008 Bergen, Norway; Department of Molecular and Cellular Biology, Harvard University, 7 Divinity Ave. Cambridge, MA 02138, USA; Department for Molecular Biology and Genetics, Cornell University, 526 Campus Road, Ithaca, NY 14853, USA; Department of Molecular & Cell Biology, Institute for Systems Genomics, University of Connecticut, Storrs, CT 06269, USA

**Keywords:** bidirectional transcription, transcription initiation, RNA decay, exosome, ncRNA, eRNA, PROMPT

## Abstract

Mammalian genomes are pervasively transcribed, yielding a complex transcriptome with high variability in composition and cellular abundance. While recent efforts have identified thousands of new long non-coding (lnc) RNAs and demonstrated a complex transcriptional repertoire produced by protein-coding (pc) genes, limited progress has been made in distinguishing functional RNA from spurious transcription events. This is partly due to present RNA classification, which is typically based on technical rather than biochemical criteria. Here we devise a strategy to systematically categorize human RNAs by their sensitivity to the ribonucleolytic RNA exosome complex and by the nature of their transcription initiation. These measures are surprisingly effective at correctly classifying annotated transcripts, including lncRNAs of known function. The approach also identifies uncharacterized stable lncRNAs, hidden among a vast majority of unstable transcripts. The predictive power of the approach promises to streamline the functional analysis of known and novel RNAs.

## INTRODUCTION

An estimated ∼75% of mammalian DNA yields RNA, at least when considering multiple cell lines^1–4^. In human cells, only ∼50% of this material is accounted for by pre-mRNA and conventional stable RNA (tRNA, rRNA, sn/snoRNA); the remaining part constitutes a population of poorly characterized lncRNA species^5^. These are mainly cell type-restricted^2^, suggesting that unknown regulatory RNAs may be found in this population. In particular the intergenic (or intervening) lncRNAs (lincRNAs) have attracted attention due to the successful functional characterization of a limited number of molecules (for recent reviews see refs.^6–10^). Other lncRNAs include promoter upstream transcripts (PROMPTs), originating in antisense orientation from active pc gene promoters^11–14^ and RNAs produced from active enhancers^14–16^ (eRNAs).

Characterization of PROMPT and eRNA production has revealed that human pc gene promoters and enhancers can be divergently transcribed^11,14,15,17–19^. A strand-bias in transcriptional directionality of pc gene promoters is apparent when considering stable RNA levels (*i.e.* seemingly producing robust amounts of mRNA in the sense direction and only little antisense PROMPT). This bias is established post-transcriptionally and governed by a decreased occurrence and utilization of early polyadenylation (pA) sites in the sense (mRNA) direction^12,13^. Such promoter-proximal pA sites trigger transcription termination and rapid transcript turnover by the ribo-nucleolytic RNA exosome complex^13^. In general, many lncRNAs are suppressed post-transcriptionally by this mechanism^20^, considerably skewing their steady-state levels from what would be expected based on transcription initiation rates alone. Therefore, transcription units that are under evolutionary pressure to evade such termination and RNA decay will constitute prime candidates for producing functional lncRNAs, which require a certain copy number for their actions.

Here we classify DNase Hypersensitive Sites (DHSs), from which capped RNA species arise in HeLa cells, by their transcriptional directionality as well as the exosome-sensitivity and abundance of their emitted RNAs. We identify stable lncRNAs with the potential to function *in trans*, unstable RNAs emanating from enhancers and a population of annotated alternative promoters, which produce exosome-sensitive mRNAs. We project that this strategy and resource of classified promoters and associated RNAs will guide annotation of functional candidates among novel and known transcripts.

## RESULTS

### Most initiation events occur divergently from DNA hotspots

With the final aim to employ biochemical criteria to characterize transcripts genome-wide, we first tested whether DHSs could be used as annotation-unbiased foci to identify transcription initiation events. Indeed, we found that ∼93% of all 5’ends of capped RNAs detected by Cap Analysis of Gene Expression (CAGE^21^) in HeLa cells^13^ were proximal to ENCODE-defined DHSs from the same cell line^22^ (Supplementary Fig. 1). The few CAGE tags not mapping to DHSs were often singletons (28% compared to 3% of DHS-proximal CAGE tags). These might be i) located within DHSs that fall beneath the peak calling cutoff for hypersensitivity, ii) represent cryptic initiation sites, iii) technical noise or iv) recapping events^23^ (although only 2% of all CAGE tags were distal to DHSs and resided in internal exons).

While most CAGE tags map to DHSs, only a small subset of DHSs initiate transcription: 58% had no proximal CAGE tags at all, and 1% and 13% of DHSs accounted for 50% and 90% of the CAGE tags, respectively (Supplementary Fig. 1). We refer to such DHSs with overlapping CAGE tags as “transcribed DHSs”. Strikingly, ∼66% (12,763 of 19,224) of transcribed DHSs showed evidence of bidirectional transcription when assessing CAGE tags derived from HeLa cells depleted of the hRRP40 (EXOSC3) exosome core component, compared to only ∼35% (6,724) DHSs from control cells (see Supplementary Fig. 2 and Methods for details). Thus, bidirectional transcription initiation is a general feature of transcribed DHSs with post-transcriptional RNA decay often affecting one strand over the other. Indeed, a large fraction (∼78%) of DHSs had at least 90% of total control CAGE expression deriving from one strand. We will refer to the dominating direction of transcription from a DHS as the ‘major’ strand and the reverse direction as the ‘minor’. Hence, unidirectionally biased transcribed DHSs generally produced largely exosome-insensitive transcripts on the major strand and exosome-sensitive transcripts on the minor strand (Fig. 1a). In contrast, RNAs produced from bidirectionally-balanced transcribed DHSs (1,986 of DHSs having at most 75% of control CAGE expression from the major strand) were generally subject to degradation by the exosome on both strands. CAGE data derived from HeLa cells depleted of hMTR4 (SKIV2L2)^19^, a nuclear-specific cofactor of the exosome, supported these observations (Supplementary Fig. 3).

**Figure 1:**
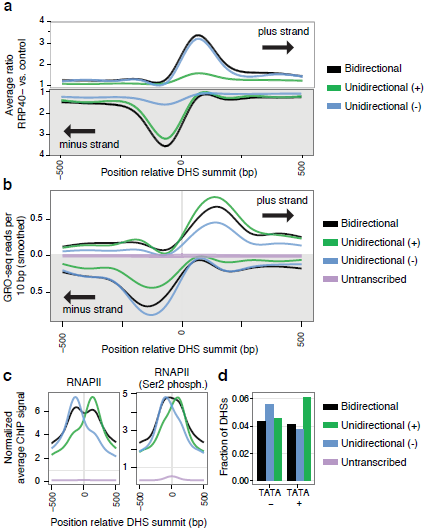
Transcriptional directionality and exosome sensitivity of RNAs emitted from DHSs. **a,** Average exosome sensitivity (vertical axis) of RNAs emanating from transcribed DHSs, broken up by strand as well as by transcriptional direction bias in control HeLa cell CAGE (indicated by color coding). Exosome sensitivity was measured as the ratio between CAGE expressions from hRRP40-depleted vs. control HeLa cells and plotted relative to DHS summits (horizontal axis). **b,** Average HeLa GRO-seq signal (per 10 bp) in a 1 kb window centered at DHS summits broken up as above. **c.** Average ENCODE^27^ HeLa ChIP signal of the RNAPII subunit POLR2A (left panel) or RNAPII with the Ser2 carboxy-terminal domain phosphorylated (right panel) in 1 kb regions centered at DHS summits, broken up as above. **d**, Fraction (vertical axis) of bidirectional and unidirectional DHSs (broken up by strand) as determined by GRO-seq data, with TATA box motif^25^ on indicated strand (horizontal axis). Average profiles in panels a-c have been smoothed.

To assay transcriptional directionality using measures which were not RNA-based, we assessed RNA polymerase II (RNAPII) initiation levels by global run-on sequencing (GRO-seq^17^) on isolated HeLa cell nuclei. TSS-proximal GRO-seq signal primarily reflects run-on activity of promoter-paused RNAPII, which is detectable as a double peak flanking the transcribed DHS. Indeed, GRO-seq data supported divergent transcriptional activity from as many as 76% (14,658) of transcribed DHSs and indicated a higher fraction (42%) of more bidirectionally balanced DHSs (Fig. 1b). We note that unidirectional DHSs (as defined by control CAGE data) on average harbored more GRO-seq signal on one strand. Thus, even though many transcribed DHSs are divergently transcribed, there may be some preference for transcription initiation in one direction, which is supported by a bias in localization and elongation status (Serine 2 phosphorylation) of RNAPII^24^ according to directionality and strand (Fig. 1c). Finally, a directional preference in transcription initiation (as measured by GRO-seq) was correlated to the presence of TATA box sites^25^ (Fig. 1d), consistent with observations in *Drosophila*^26^.

Based on the above analyses, we conclude that transcription is typically initiated in both directions from a limited number of accessible DNA hotspots. Taken together with the observed strand-specific bias in exosome sensitivity, this implies that the previously characterized properties of certain promoter/DHS subclasses (mRNA-PROMPT pairs^13^ and eRNAs^19^) are general features of regions that initiate transcription.

### Directional bias and exosome sensitivity discern RNA biotypes

Having established that bidirectionality is general for transcribed DHSs, we next assessed whether this and exosome sensitivity of the produced transcripts could be used at a broader scale for RNA species classification. We employed GENCODEv17^5^ to subdivide a set of transcribed DHSs (as observed after hRRP40 depletion) into 9,040 mRNA- and 637 lncRNA-promoters with no ambiguous annotation as well as 12,731 DHSs with no annotation support (Supplementary Table 1-3, see Methods). As expected, mRNAs were not, or only mildly, exosome-sensitive (Fig. 2a), while the clear majority of their antisense PROMPTs were (Fig. 2b). Moreover, most transcripts originating from unannotated DHSs displayed strong exosome sensitivity on both strands. Annotated lncRNAs fell between these two extremes with the majority being exosome sensitive. Finally, and consistent with previous studies^2,13^, the steady state abundance, measured in control CAGE samples, was much higher (average ∼17-fold) for mRNAs than for unannotated transcripts, again leaving lncRNA abundance in between the two (Fig. 2c).

**Figure 2:**
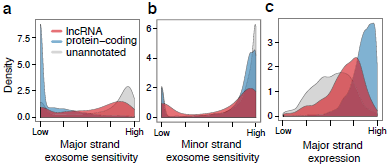
Exosome sensitivities and expression levels of annotated transcript biotypes. **a**, Densities of exosome sensitivity of RNAs emanating in the major direction (defined as most dominating in the control CAGE library) of considered DHSs, broken up by GENCODEv17 TSS annotations associated with the corresponding DHSs. Sensitivity was defined as the relative fraction of measured RNA abundance reduced by the exosome (the fraction of hRRP40 CAGE expression level not detected with control CAGE, see Methods). **b,** As panel **a**, but assessing RNAs on the opposite (minor) strand. **c**, Expression levels from DHSs, associated with the indicated RNA biotypes, assessed as ranked CAGE expression levels in control HeLa cells of RNAs emanating from major strands at DHSs broken up by annotation as above.

The obvious separation of the three RNA biotypes based on such simple criteria prompted us to test whether the same biological features could be used to broadly distinguish core promoters of RNA species. To this end, we considered properties describing RNAs emitted from their respective DHSs: their overall and strand-specific expression levels, their strand bias (directionality) and their overall and strand-specific sensitivity to the exosome (Methods). Gratifyingly, 80% of the total variance in these seven dimensions could be explained by only two principal components (Fig. 3a). We employed *k*-medoids clustering to discern five major DHS groups based on the same properties (Supplementary Table 1). The resulting clusters showed distinct patterns of expression and exosome sensitivity (Fig. 3a-b). DHSs from the five clusters expressed RNAs with distinct enrichments and depletions of GENCODEv17 annotated transcript biotypes (Fig. 3c and Supplementary Fig. 4a, see Methods). Specifically, the two clusters of *unidirectional stable* DHSs (Fig. 3a-b, red and blue) were highly enriched for mRNA TSSs with unannotated (Fig. 3c, light blue column, odds ratio (OR)=34.5) or annotated (Fig. 3c, orange column, OR=10.5) minor strand lncRNA neighbors. Since these two clusters are strand-specific mirrors of one another, they were merged into a single class for the remaining analyses. Major strand RNAs originating from these DHSs were abundant in control CAGE samples, while their corresponding minor strand transcripts were lowly transcribed and highly exosome-sensitive (Fig. 3b). This is reminiscent of PROMPT-mRNA transcript pairs^13^ and likely reflects that moderately abundant PROMPTs have previously been annotated as lncRNAs, while lowly abundant ones have not been annotated at all. Conversely, the *intermediate* and *weak unstable* DHS clusters (Fig. 3a-b, orange and green) were strongly depleted of mRNA TSSs (Fig. 3c, orange, light blue and blue bars, combined OR=0.26). In contrast, both were enriched for unannotated TSSs (OR=14.2 for *intermediate unstable* and OR=105.7 for *weak unstable*). The former cluster was also enriched for annotated lncRNAs (OR=5.4, red bar). DHSs in these two clusters had a more balanced bidirectional expression with stronger exosome sensitivity on both strands, when compared to *unidirectional stable* DHSs (Fig. 3b). We note that the two *unstable* clusters in reality form a gradient, with increasing expression and decreasing exosome sensitivity from *weak unstable* DHSs to *intermediate unstable* DHSs. Finally, the *bidirectional stable* DHS cluster (Fig. 3a-b, purple) was enriched for bidirectionally transcribed and exosome-insensitive pc transcript-derived RNAs (OR=49.1). Annotated lncRNAs were also enriched in this cluster as well as in both the *unidirectional stable* and *intermediate unstable* DHS clusters, reflecting the heterogeneity of lncRNAs in terms of abundance and exosome sensitivity (Fig. 3b,c).

**Figure 3:**
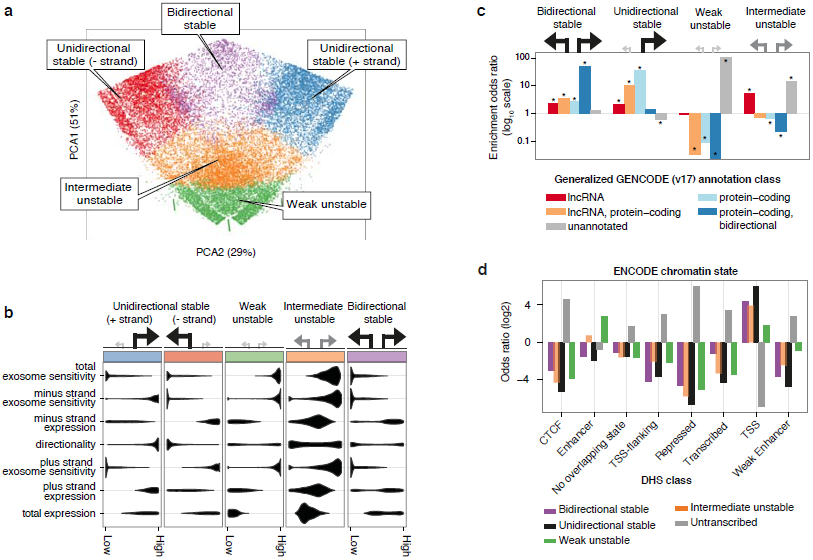
Transcriptional directionality, RNA abundance and exosome sensitivity separate functionally distinct groups of promoters. **a**, Transcribed ENCODE HeLa DHSs were grouped into five major classes via *k*-medoids clustering based upon exosome sensitivity, expression levels and transcriptional strand bias (directionality). DHSs and their cluster memberships were visualized by principal component analysis (PCA). The first two principal components describe ∼80% of the total variance in the data used for clustering. Colors and names given to each DHS class are utilized throughout the paper. **b**, Biochemical properties of each DHS cluster are summarized by horizontal density plots. Illustrations on top of subpanels depict the typical arrangement of TSSs within each DHS cluster where the size and shade of the arrows indicate abundance and stability of emitted RNAs, respectively. The two unidirectional stable clusters are merged in subsequent analyses. **c**, Enrichment odds ratios (vertical axis, log_10_ scale) of DHS overlap with TSSs of GENCODEv17 transcripts (annotation classes are described in Methods). Stars indicate Fisher’s exact test Benjamini-Hochberg *FDR* < 0.001. **d,** Enrichment odds ratios (log_2_-transformed, vertical axis) of DHS overlap with ENCODE chromatin segmentation states. While all clusters of transcribed DHSs are enriched for predicted promoters in HeLa cells (‘TSS’; OR ranging between 3.6 and 61.4), weak unstable DHSs are highly enriched for predicted strong enhancers (‘E’; OR=6.9). Note that untranscribed DHSs mainly fall into the classifications not associated with predicted active transcription initiation (‘repressed’, ‘CTCF’, ‘transcribed’, and ‘promoter-flanking’ states).

To characterize the DHSs that produce unannotated RNAs, we investigated the chromatin status of DHSs using ENCODE HeLa chromatin segmentation states^27^ (Supplementary Table 4). While all DHS clusters were enriched for predicted gene promoter chromatin states (‘TSS’, OR ranges from 3.6 to 61.4), *weak unstable* DHSs were highly enriched for chromatin-predicted enhancers (OR=6.9, ∼40% overlap vs. < 4% overlap with stable DHSs) (Fig. 3d and Supplementary Fig. 4b). Furthermore, this cluster contained the highest enrichment for ChIP-seq signals of enhancer-associated histone modification (H3K4me1) and proteins, including *FOS/JUN*, *P300* and the cohesin component, *SMC3* (Supplementary Fig. 5). This strongly indicates that weak unstable DHSs to a large extent represent transcribed enhancers.

In summary, the overlap analyses of the DHS clusters with gene annotations and chromatin states independently confirm that the CAGE-based discrimination approach captures biochemically distinct properties of transcribed DHSs in terms of their produced transcripts.

### DHS clusters reveal distinct RNA properties and constraints

To investigate the properties of RNA produced from the clustered DHSs without relying on annotation, we assembled *de novo* transcripts^28^ from control and hRRP40-depleted RNA-seq libraries previously obtained from HeLa cells^13,29^ (Supplementary Table 5). Association of assembled transcripts with the classified DHSs (Supplementary Table 6) revealed several interesting relationships. The *bidirectional stable* and the major strand of the *unidirectional stable* DHSs generally produced multi-exonic transcripts of >2000 nt (Fig. 4a and Supplementary Fig. 6a). Conversely, mostly mono-exonic and shorter transcripts (around 1000 nt or shorter) derived from the minor strand of *unidirectional stable* DHSs and from both strands of *unstable* DHSs. Thus, RNAs originating on the minor strand from e.g. pc gene promoters (PROMPTs) display similar structures to transcripts from unstable DHSs. Consistent with their stable and multi-exonic nature (Fig. 4a), the majority (∼83%) of RNAs emanating from *stable* DHSs are likely protein-coding as estimated by PhyloCSF^30^ (Supplementary Fig. 6b and Supplementary Table 7, also see Methods). In contrast, 87-97% of RNAs from *unstable* DHSs or PROMPTs are likely non-coding. Interestingly, the small fraction of unstable RNAs with protein-coding potential constitute transcripts that typically also are different in terms of transcript structure (i.e. number of exons and length, see Supplementary Fig. 6c-d). Corroborating these results, we note that unstable RNAs were to a lesser extent polyadenylated and more nuclear-retained than those derived from stable DHSs (Supplementary Fig. 7), based on ENCODE CAGE and RNA-seq fractionation data^2^.

**Figure 4:**
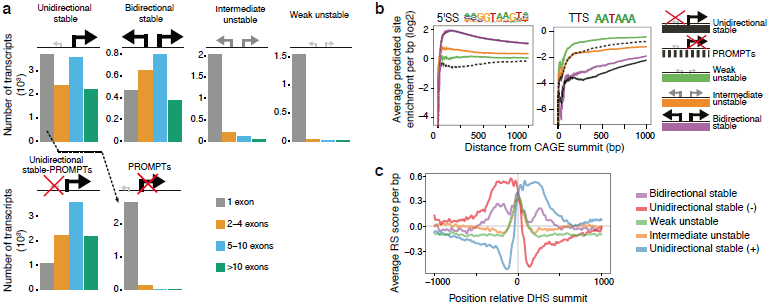
RNA processing events separate DHS clusters. **a**, Histograms of *de novo-*assembled transcript counts, broken up by number of exons and associated DHS cluster. Vertical axes indicate the number of thousand transcripts. In the bottom panels, the unidirectional stable DHSs are split on major (left) and minor (right) strands, revealing that PROMPTs are highly similar to transcripts of weak unstable DHSs. **b**, Frequencies of RNA processing motifs (5’SS, left panel, and pA-site hexamer, right panel) downstream (major strand) of CAGE summits broken up by DHS cluster. Vertical axis shows the average number of predicted sites per kb within an increasing window size from the TSS (horizontal axis) in which the motif search was done. 0 indicates the expected hit density from random genomic background. **c**, Average number of rejected substitutions across mammals per bp around summits of transcribed DHSs, broken up by DHS class.

In line with their low intron content, the prevalence of 5′ splice site (5'SS) motifs was at, or near, genomic background levels downstream of the TSSs of *unstable* DHSs and minor strand TSSs of *unidirectional stable* DHSs (Fig. 4b, left panel). Conversely, 5'SS motifs were highly over-represented downstream of *bidirectional* and major strand TSSs of *unidirectional stable* DHSs. Consistent with earlier comparisons of motif occurrences downstream of PROMPT-, eRNA- and mRNA-TSSs^12,13,19^ we found that the 5'SS motif frequency was anti-correlated with the downstream frequency of proximal consensus pA site hexamer AAUAAA motifs (Fig. 4b). The 5'SS motifs prevent the utilization of TSS-proximal pA sites^31^, which otherwise leads to exosomal decay^13,19^. Therefore, our observation that pA sites are generally depleted in TSS-proximal regions downstream of stable RNAs compared to regions flanking *unstable* DHSs as well as downstream of PROMPT TSSs (Fig. 4b, right panel) provides a mechanistic explanation for the observed differences in RNA stability between the defined DHS clusters.

Given that pA- and 5'SS frequencies downstream of TSSs are highly related to RNA stability, these features are perhaps under selective pressure to ensure the proper production and stability of functionally relevant RNAs. Indeed, we generally found exosome-insensitive RNAs to be produced from evolutionary constrained DNA (Fig. 4c). In contrast, PROMPTs and unstable RNAs were produced from DNA with a notably faster evolutionary rate. In other words, unbalanced evolutionary rates in DHS-flanking regions are highly predictive of transcriptional strand bias (Supplementary Fig. 8). While the core DHS region is evolutionarily constrained regardless of its class, the average number of rejected substitutions (RS) between mammals^32^ in regions flanking unstable DHSs is lower than expected, indicating selectively rapid evolution, as also previously noted^33^. Weak unstable DHSs thus bear a resemblance to transcribed enhancers identified previously from lncRNA-associated DHSs with enhancer-characteristic chromatin marks ^34^.

Taken together, these analyses demonstrate that classification of TSSs by the nature of the RNAs they emit in a bidirectional pattern is not only predictive of annotated RNA biotypes but also reflects associated properties: RNA lengths, 3’end processing and splicing events, protein-coding content, cellular localization and evolutionary constraints.

### Characterization of RNAs from known and novel promoters

Having established the predictive power of our approach, we systematically classified 24,007 transcribed HeLa DHSs, which were either unannotated or associated with GENCODE annotated TSSs of mRNAs or lncRNAs (thereby not considering TSSs of, for instance, annotated pseudogenes and short RNAs; see Methods). We associated these with *de novo* derived RNA-seq transcripts with the aim to characterize novel TSSs and to identify outliers within annotated transcripts classes. The classification strategy, outlined in Fig. 5a, shows the number of transcribed DHSs passing each filtering step according to both lenient (only DHS cluster association) and strict (additional exosome sensitivity threshold) criteria. The classification showed that while *unstable* DHSs typically produced unannotated RNAs, 3,046 (23%) of stable DHSs were, surprisingly, not associated with any TSS annotation. RNAs from these DHSs were highly enriched in 5’UTRs, and to a lesser extent in exons and introns of pc gene models (Fig. 5b). Therefore, they likely derive from yet-to-be annotated alternative mRNA promoters. In fact 654 (21%) of these DHSs produced major strand transcripts detected by RNA-seq, and 353 of these had RNA-seq-derived exons overlapping with annotated exons (Supplementary Table 8). An example of this is a DHS corresponding to a novel TSS responsible for a large majority of mRNA production (as measured by control CAGE and RNA-seq) from the *TULP4* gene (Fig. 6a, blue shadowed region).

**Figure 5.**
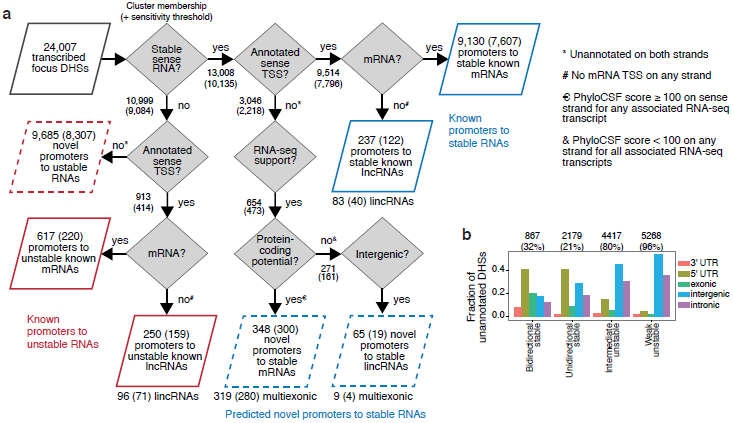
RNA annotation by means of DHS classification. **a.** Flow chart illustrating the filtering steps made to extract transcribed DHSs with interesting properties, such as those emitting unstable mRNAs or novel stable multi-exonic lincRNAs. The number of DHSs passing each filtering step based on lenient (DHS cluster membership) and strict (additional sensitivity thresholding) criteria is indicated at each arrow. For stable and unstable DHSs an exosome sensitivity threshold of ≤ 0.25 and ≥ 0.75, respectively, was used in addition to DHS cluster membership for strict filtering. The subset of 24,007 transcribed HeLa DHSs, which were either unannotated or associated with GENCODE annotated TSSs of mRNAs or lncRNAs (see Methods) were considered. **b.** Fraction of unannotated DHSs (not overlapping known TSSs) overlapping other genomic features, broken up by DHS cluster. Absolute numbers and fractions of DHSs in each class falling into the unannotated category are shown on top.

Conversely, 617 DHSs associated with GENCODEv17 annotated mRNA TSSs (corresponding to 609 genes) emitted unstable RNAs (Supplementary Table 9). Strikingly, 246 (∼40%) of these were associated with genes that also produce stable RNAs from another DHS. Illustrating this, the *TGIF1* locus has three alternative promoters that produce sense transcripts with vastly different exosome sensitivity and abundance (Fig. 6b). Interestingly, when compared to stable mRNAs, the DNA downstream of these internal exosome-sensitive mRNA TSSs was not enriched in elongation chromatin marks (H3K79me2, H3K36me3, H4K20me1) (Fig. 6c and Supplementary Fig 9). We hypothesize that these TSSs are producing RNAs, which do not extend to the canonical pA site. This suggests that unstable mRNAs are in fact similar to eRNAs and PROMPTs not only in terms of their low stability, but also in terms of their early transcriptional termination. Indeed, downstream pA site frequencies separated highly unstable annotated mRNAs from highly stable ones (Supplementary Fig. 10a). However, ∼49% of these mRNA promoters were also supported by RefSeq curated mRNA TSSs (compared to ∼72% of the highly stable mRNA promoters), and had similar expression distribution across human cell types^35^ as stable mRNAs (Supplementary Figs. 10b-c). Thus, many of these DHSs produce stable, or at least reproducibly detectable, mRNAs in at least some cell types.

**Figure 6:**
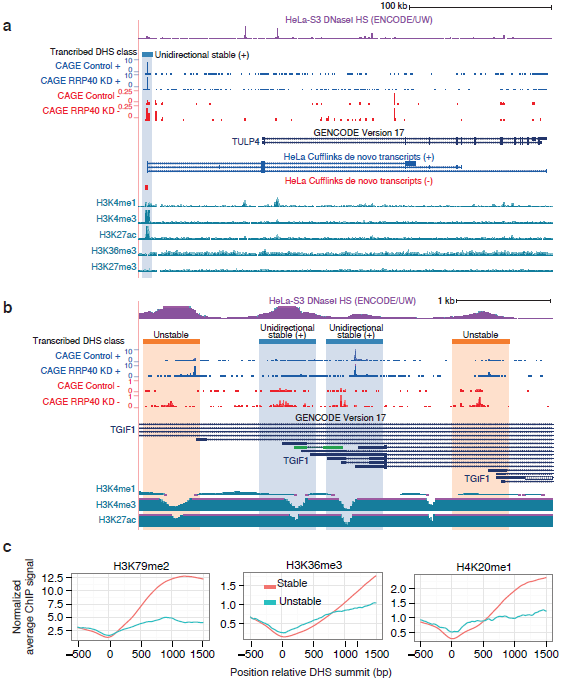
Examples of DHS class-facilitated RNA characterization. **a**, UCSC genome browser image showing a transcribed DHS corresponding to a novel promoter (chr6:158653065-158653215, hg19) in the *stable unidirectional* class, linked to the *TULP4* gene by RNA-seq and CAGE (blue highlight). This promoter accounts for the majority of stable *TULP4* expression in HeLa cells. Average replicate CAGE expression before and after exosome (hRRP40) depletion, and Cufflinks *de novo* assembled transcripts connecting the novel promoter with canonical exons are shown. Note the different scales for plus and minus strand CAGE. Below, ENCODE ChIP-seq profiles of H3K4me1, H3K4me3, H3K27ac, H3K36me3 and H3K27me3 are shown. **b**, UCSC genome browser image showing the classification of transcribed DHSs corresponding to alternative promoters within the *TGIF1* gene, which differ substantially in RNA output and stability. Average replicate CAGE expression before and after exosome (hRRP40) depletion and ENCODE ChIP-seq profiles of H3K4me1, H3K4me3 and H3K27ac are shown. Note that the ChIP-seq data lack the resolution to ascertain usage of promoters. **c**, Average ChIP-seq signal (vertical axis) of H3K36me3, H3K79me2 and H4K20me1 (associated to productive elongation), -500 to + 1500 bp (with respect to major strand) around summits of DHSs (horizontal axis) emanating stable and unstable mRNAs. The ChIP-seq signal was normalized to the overall signal across all ENCODE HeLa DHSs.

Among stable unannotated DHSs with intergenic RNA-seq transcript support, we found no likely mRNA candidates as concluded from their on-average low phyloCSF^30^ scores. This suggests that the vast majority of pc genes expressed in HeLa cells are already discovered. However, 9 stable DHSs marked promoters of multi-exonic lincRNAs (Supplementary Table 10), representing a small but interesting set of non-coding RNAs with putative functions *in trans*.

While annotated lincRNAs, such as *NEAT1* and *MIR17HG*, were typically exosome sensitive, there were exceptions, including *H19, FTX, TINCR, HCG11, LINC00473* and *SNHG16* (Supplementary Table 11). Some lincRNAs function as primary precursors for the production of smaller ncRNAs such as microRNAs (e.g. H19) and snoRNAs (e.g. *SNHG16*). While the endocleavage events of miRNA biogenesis make this process independent of splicing, the opposite is the case for snoRNAs, which are matured by exonucleoytic trimming after their liberation by splicing^36^. Thus, lincRNAs hosting snoRNAs are expected to have a strong evolutionary pressure to ensure proper splicing, while this is not the case for miRNA-hosting lincRNA. Indeed, snoRNA host gene lincRNAs are less exosome-sensitive than miRNA-hosting as well as non-host lincRNAs (P < 0.004, Mann-Whitney U test) (Supplementary Fig. 11). Further illustrating this, roughly 25% of exosome-insensitive lincRNA transcripts are snoRNA hosts, compared to ∼1% of exosome-sensitive lincRNA transcripts.

## DISCUSSION

In this work, we have presented a classification of DHSs based on the abundance, directionality and exosome sensitivity of their emitted transcripts. The approach, which relies on the widespread bi-directional nature of transcription initiation in human cells, efficiently distinguishes known transcript classes and their related properties, such as processing status, cellular localization, and evolutionary constraints.

We have shown that the core regions of classified DHSs are highly similar between DHS classes in terms of their conservation and bidirectional initiation events. Thus, the distinction between human enhancers and promoters is fuzzy, especially since they are also similar in terms of transcription factor binding sites^19^, core promoter-like elements^19^, and binding of general transcription factors^37^. In the same line, a subset of intragenic enhancers have been reported to work as gene promoters^38^.

In fact, the strongest feature distinguishing enhancers and promoters seem to be the characteristics of their produced RNAs, which are at least partially determined by processing motifs downstream of the respective TSSs.

While our method can be used to identify novel RNAs, its main strength is that it provides a unique inroad to characterize lncRNAs, which are only known by their transcript structures. Importantly, the approach is based on biochemical properties of transcriptionally active regions, independent of current gene annotations. For example, we identify 353 novel stable promoters that can be linked to known pc genes by RNA-seq, and 9 novel multi-exonic stable lncRNA promoters that are promising candidates for functional validation. For previously identified RNAs, we find that only few annotated lncRNAs, including lincRNAs, are resistant to exosome-mediated decay. Hence, a considerable number of ncRNAs are unlikely to be functional as high-copy molecules *in trans*, unless cell-type-specific mechanisms exist to regulate their turnover. This does not suggest that the gene models for these lncRNAs are of low quality, only that the expressed RNAs are susceptible to exosome-mediated degradation. Conversely, the large majority of transcribed mRNAs are exosome-insensitive.

Our approach enables the identification of promoters with unexpected properties that highlight important mechanistic questions to guide future studies. One example is pc genes with alternative promoters displaying differential exosome sensitivity; the most sensitive alternative promoters likely do not produce a protein-coding product. While the most exosome-sensitive alternative promoters have the hallmark pA site frequency of other unstable RNA classes, it is unclear how stable RNA produced by upstream promoters of the same gene are avoiding this fate since the same pA sites are encountered during transcription. In-depth studies of these cases might reveal mechanisms that cells use to stabilize normally unstable RNAs in given circumstances or cell types. It also suggests that stable transcripts are under continual selective pressure to avoid early pA sites and include splice sites in order to stay stable, while the default state of the human genome is to discourage the generation of long stable RNAs.

## METHODS

### HeLa cells culturing and siRNA-mediated knockdowns

HeLa cells originating from the S2 strain were cultured and transfected with EGFP (control), hMTR4 (*SKIV2L2*), or hRRP40 (*EXOSC3*) siRNA performed as described previously^13,19^. Briefly, cells cultured in DMEM/10%FBS/1%Pen/Strep were transfected using Lipofectamine 2000 (Life Technologies) and a final siRNA concentration of 20 µM. Transfections were performed 1 and 3 days after cell seeding and harvested on day 5.

The following siRNA sequences were used:

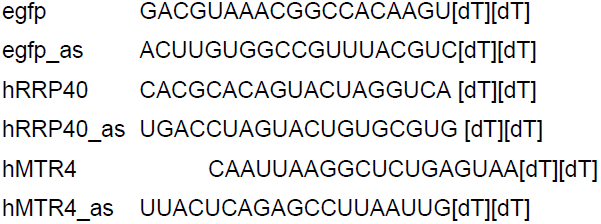

### HeLa CAGE library preparation, sequencing and mapping

Previously sequenced HeLa CAGE libraries^13^ (GEO IDs GSE48286 and GSE49834) were extended with two additional biological replicates per condition (GEO ID GSE58991). The same methods for preparation and computational processing were used as described in these reports. In brief, CAGE libraries were prepared as described in ref.^21^ from 5 μg of total RNA purified from 2×10^6^ HeLa cells using the Purelink mini kit (Ambion) with 1% 2-Mercaptoethanol (Sigma) and on-column DNAse I treatment (Ambion). Reads were trimmed to remove linker sequences and subsequently filtered for a minimum sequencing quality of 30 in 50% of the bases. Mapping to the human genome (hg19) was performed using Bowtie^39^ (version 0.12.7), allowing for multiple good alignments and subsequently filtering for uniquely mapping reads.

### GRO-seq library preparation and processing

Libraries were prepared as in ref.^26^, using the adapter ligation protocol and starting with at least 5×10^6^ HeLa nuclei. Briefly, a ribonucleotide analog [5-bromouridine 5′-triphosphate (BrUTP)] was added to BrU-tag nascent RNA during the run-on step. Nuclear Run-on (NRO) RNA was chemically hydrolyzed into short fragments (∼100 bases). BrU-containing NRO-RNA was triple-selected through immunopurification, in parallel with RNA end repair and adapter ligations. NRO-cDNA libraries were then prepared for sequencing. The libraries were sequenced on the Illumina HiSeq2500, using standard protocol at the Cornell bioresources center (www.BRC.cornell.edu). Reads were trimmed to 30 bases and first mapped to a representative complete transcribed unit of rDNA (GenBank accession id: U13369.1) using Bowtie^39^. Reads not aligning to the rDNA were then mapped to the human genome (hg19). Reads that mapped uniquely with 2 mismatches or less were then used for downstream analyses.

### DHSs as focus points for transcription initiation

For unbiased categorization of TSSs, we focused on ENCODE HeLa DNase I hypersensitive sites^22^ (DHSs). A set of 199,188 combined (UW and Duke) FDR 1% peaks (narrowPeaks, ENCODE Jan 2011 integration data) were considered. For each DHS we required a well-defined DNase signal summit (position with max DNase signal), hereafter referred to as DHS summit, supported by either UW or Duke DNase data (Jan 2011 ENCODE integration data). Immediately flanking these DHS summits, we defined two windows of size 300 bp associated with minus and plus strand expression as illustrated in Supplementary Fig. 2. We further filtered DHSs to not overlap any other DHS strand-specifically with respect to these windows. This resulted in a set of 178,655 genomic well-separated DHSs with well-defined DHS summits.

### Quantification of DHS-associated expression

DHS-associated strand-specific expression in control and exosome (hRRP40) depleted HeLa cells were quantified by counting of CAGE tags in genomic windows of 300 bp immediately flanking DHS summits (as described above). CAGE tag counts were then converted to tags per million mapped reads (TPMs). After inspection of preferential location of CAGE tags with respect to strand around DHS summits (not shown), we decided to focus on transcription going outwards from the DHS summits. Hence, unidirectional and divergent but not convergent transcription was considered. 81% of all CAGE tags were covered by the filtered set of DHSs and these flanking windows. For subsequent analyses, we required DHSs to be supported by CAGE tag start sites (CTSSs) of at least 2 CAGE tags on the same strand in at least two replicates and with an average replicate expression level of at least 0.5 TPM. This resulted in 19,224 and 25,342 transcribed DHSs in control HeLa cells and in HeLa cells after exosome (hRRP40) depletion, respectively. For estimation of the number of divergently transcribed DHSs, we counted the number of DHS with CTSSs supported by at least 2 CAGE tags on both strands in at least one replicate CAGE library.

### Measuring directionality of transcription

Based upon strand-specific DHS expression (described above), we calculated a directionality score measuring the strand bias in expression level for transcribed DHSs. The directionality score (*D*) measures the transcriptional bias to either plus (*P*) or minus (*M*) strand of each DHS:

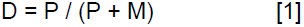

*D* ranges between 0 (100% minus strand expression) and 1 (100% plus strand expression, and 0.5 indicates a perfectly balanced bidirectional output. DHSs with a directionality score ≤ 0.1 or above ≥ 0.9 were considered ‘unidirectional’ biased while DHSs with a directionality score ≥ 0.25 and ≤ 0.75 were considered ‘bidirectional’.

### GRO-seq based analysis of CAGE-expressed DHSs

We used GRO-seq to estimate the directionality and the frequency of bidirectional transcription initiation of transcribed DHSs (as defined by CAGE) in the same way as described above for CAGE data (but using GRO-seq reads instead of CAGE tags). A strand-specific search with a TATA box motif^25^ was done in 601 bp regions focused on DHS summits using the ASAP tool^40^ (with standard settings), and with 0.9 relative score as a cutoff. The frequency of predicted TATA sites on each strand was then calculated for unidirectionally biased or bidirectionally balanced DHSs (as determined from GRO-seq data).

### Measuring exosome sensitivity

Based upon strand-specific expression (described above) we calculated a strand-specific exosome sensitivity score measuring the relative amount of degraded RNAs by the exosome. We designed the sensitivity score to quantify the fraction of hRRP40 depleted CAGE expression seen only after exosome depletion. Exosome sensitivity was calculated for both strands:

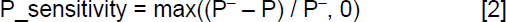

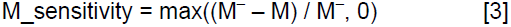

*M^−^* and *P^−^* denote the expression level in minus and plus strand windows after exosome (hRRP40) depletion, respectively, while *M* and *P* denote the expression level in minus and plus strand windows in control HeLa cells. Similarly, we calculated an overall exosome sensitivity score for each transcribed DHS after first adding plus and minus strand expression.

For specific analyses, we used thresholds of ≤ 0.25 and ≥ 0.75 to identify highly stable and highly unstable RNAs emanating from transcribed DHSs.

### GENCODE transcript-association with DHSs

We annotated DHSs with GENCODE^5^ version 17 transcripts. Each gene transcript with an annotated TSS overlapping a DHS window of the same strand (described above) was associated with that DHS. We further generalized the GENCODE gene_type:transcript_type biotypes according to the scheme in Supplementary Table 12. Each DHS was assigned one generalized biotype using the hierarchical strategy combining the generalized biotypes associated with each strand (Supplementary Table 13). For each DHS generalized biotype, both the minus and plus strand criteria had to be fulfilled by at least one associated transcript. DHSs that were assigned a generalized biotype were not considered in lower ranked tests. Analyses specifically considering lincRNAs were based on lncRNA DHSs associated with at least one GENCODE transcript of transcript_type lincRNA.

### *k*-medoids clustering of DHSs

*k*-medoids clustering was performed *k*=2-10 clusters on DHSs with observed expression after exosome (hRRP40) depletion (see calculation of expression above), based on DHS-associated exosome sensitivity on minus and plus strands, overall exosome sensitivity, directionality of transcription after exosome (hRRP40) depletion, total HeLa control expression as well as HeLa control expression for minus and plus strands. Expression levels were converted to ranks and rescaled to [0,1]. This data were used for visualization of DHSs by principal component analysis (PCA). Before *k*-medoids clustering of DHSs, all data were centered (values had their means subtracted) and rescaled (values were divided by their standard deviations). The final number of clusters (*k*=5) was selected according to clustering purity and entropy^41^ with respect to DHS-associated generalized GENCODE biotypes, based on a local maxima with no further increase in purity and only a marginal decrease in entropy (2.6%).

### ENCODE segmentation state association with DHSs

DHSs were categorized into various chromatin states according to overlap of DHS summits with combined Segway^42^ and ChromHMM^43^ ENCODE (release Jan 2011) HeLa state segmentations.

### *De novo* assembly of transcripts from RNA-sequencing data

Transcripts were assembled *de novo* from RNAseq data from HeLa cells depleted of hRRP40^29^ (SRA accession: SRX365673). We utilized Cufflinks v2.1.1^28^ applying standard parameters with the following modification to enhance assembly of low-abundant transcripts: --*min-frags-per-transfrag 5* (only require 5 read fragments to assemble a transcript) and --*overlap-radius 150* (allow gaps of up to 150 bp between fragments assembled into the same transcript). Assembled transcriptomes were then converted to bed format and paired with DHS regions by overlap with transcript 5’ ends using BEDTools^44^. Transcript length and exon count was then extracted from the DHS-associated set of *de novo* assembled transcripts.

### Poly-adenylation status of RNA-seq derived transcripts

ENCODE RNAseq libraries from polyA+ and polyA-fractions (SRA accessions: SRX084680 and SRX085297, respectively) were mapped to all DHS-associated *de novo-*assembled transcripts using the STAR pipeline^45^ and RPKM values were calculated using the rpkmforgenes.py script^46^.

### Protein-coding **p**otential of *de novo*-assembled transcripts

PhyloCSF^30^ was used to evaluate the coding potential of Cufflinks *de novo* assembled transcripts whose TSSs were associated with transcribed DHSs. PhyloCSF was run in start-to-stop ORF mode (ATGStop) on alignments from 29 mammals (http://hgdownload.soe.ucsc.edu/goldenPath/hg19/multiz100way/maf/). Transcripts were divided into two groups based on protein-coding potential: predicted proteins (PhyloCSF score ≥ 100) and predicted non-coding RNAs (PhyloCSF score < 100). The threshold was selected based on results from ref. ^47^.

### Analysis of downstream splice site and termination signals

To investigate the preference of RNA processing motifs downstream of TSSs potentially differing between unstable and stable DHSs, we first identified the genomic distributions of the splice site and AATAAA termination motifs (motifs determined elsewhere^19^) using HOMER^48^. We then calculated the enrichment per bp of these motifs, in regions of increasing sizes (up to 1kb) downstream of the CAGE summit (position with maximum signal over pooled CAGE libraries) associated with respective strand of each DHS, compared to the expected number of motifs according to a uniform genomic distribution.

### Chromatin state and transcription factor binding at DHSs

ENCODE HeLa ChIP-seq pileup data (release Jan 2011) were extracted around DHSs, averaged per base pair with respect to distance to DHS summits and DHS cluster/category and normalized by division with the average overall pileup signal around all DHSs. Hence, a normalized signal >1 indicates more signal than would be expected by chance around DHSs and a normalized signal <1 indicates less signal than would be expected by chance around DHSs.

### Analysis of DHSs with FANTOM5 CAGE data

We quantified the expression of transcribed DHSs, as described above, using FANTOM5 primary cell CAGE data^35^ For each DHS and strand, both the max expression and a cell type-specificity score were calculated. The specificity score was defined to range between 0 and 1, where 0 means unspecific (ubiquitously expressed across cell types) and 1 means specific (exclusively expressed in one CAGE library). In detail,

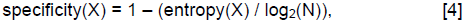

where *X* is a vector of expression values for a DHS over all CAGE libraries and *N* its cardinality *(|X|*, the number of CAGE libraries).

### Evolutionary conservation

Evolutionary constraints on DNA surrounding DHS summits were estimated using Genomic Evolutionary Rate Profiling (GERP) data (http://mendel.stanford.edu/SidowLab/downloads/gerp/). We used the available estimated Rejected Substitutions (RS) across mammalian alignments as an indicator of the strength of past purifying selection and deviance from neutral evolutionary rate.

### Statistics and visualization

Statistical tests were done in the R environment (http://www.R-project.org). Graphs were made using mainly ggplot2 R package. Intersections of and distances between various genomic features were calculated using BEDTools^44^.

## Acknowledgements

This work was supported by grants from NIH (grant GM025232 supporting L.C.), the Novo Nordisk and Lundbeck foundations (to A.S.) and the Danish National Research Foundation (grant DNRF58), the Danish Cancer Society and the Lundbeck- and Novo Nordisk Foundations (to T.H.J.).

### Author contributions

RA, PRA, EV and AS made the computational analyses. LJC made the GRO-seq libraries. JB and MB made the CAGE libraries. THJ and AS supervised the project. RA, PRA, THJ and AS wrote the paper with input from all authors.

### Competing Financial Interests

The authors declare no competing interests

### Data accessibility

CAGE and GROseq data generated for this study were deposited in the GEO database (ID:GSE58991). The CAGE libraries expanded our previous experiments, which were also used in this study (GEO IDs: GSE48286 and GSE49834). Similarly, RNA-seq data used in the study were previously deposited in the Short Read Archive (ID: SRX365673)

## REFERENCES

1. Birney, E. et al. Identification and analysis of functional elements in 1% of the human genome by the ENCODE pilot project. Nature 447, 799–816 (2007).

2. Djebali, S. et al. Landscape of transcription in human cells. Nature 489, 101–108 (2012).

3. Kapranov, P. et al. RNA maps reveal new RNA classes and a possible function for pervasive transcription. Science 316, 1484–1488 (2007).

4. The FANTOM Consortium. The Transcriptional Landscape of the Mammalian Genome. Science 309, 1559–1563 (2005).

5. Derrien, T. et al. The GENCODE v7 catalog of human long noncoding RNAs: analysis of their gene structure, evolution, and expression. Genome Research 22, 1775–1789 (2012).

6. Rinn, J. L. & Chang, H. Y. Genome regulation by long noncoding RNAs. Annual review of biochemistry 81, 145–166 (2011).

7. Ulitsky, I. & Bartel, D. P. lincRNAs: Genomics, Evolution, and Mechanisms. Cell 154, 26–46 (2013).

8. Lam, M. T. Y., Li, W., Rosenfeld, M. G. & Glass, C. K. Enhancer RNAs and regulated transcriptional programs. Trends in Biochemical Sciences 39, 170–182 (2014).

9. Jensen, T. H., Jacquier, A. & Libri, D. Dealing with Pervasive Transcription. Molecular Cell 52, 473–484 (2013).

10. Cech, T. R. & Steitz, J. A. The Noncoding RNA Revolution—Trashing Old Rules to Forge New Ones. Cell 157, 77–94 (2014).

11. Preker, P. et al. RNA Exosome Depletion Reveals Transcription Upstream of Active Human Promoters. Science 322, 1851–1854 (2008).

12. Almada, A. E., Wu, X., Kriz, A. J., Burge, C. B. & Sharp, P. A. Promoter directionality is controlled by U1 snRNP and polyadenylation signals. Nature 499, 360–363 (2013).

13. Ntini, E. et al. Polyadenylation site-induced decay of upstream transcripts enforces promoter directionality. Nat Struct Mol Biol 20, 923–928 (2013).

14. Sigova, A. A. et al. Divergent transcription of long noncoding RNA/mRNA gene pairs in embryonic stem cells. Proceedings of the National Academy of Sciences 110, 2876–2881 (2013).

15. Kim, T.-K. et al. Widespread transcription at neuronal activity-regulated enhancers. Nature 465, 182–187 (2010).

16. De Santa, F. et al. A Large Fraction of Extragenic RNA Pol II Transcription Sites Overlap Enhancers. Plos Biol 8, e1000384 (2010).

17. Core, L. J., Waterfall, J. J. & Lis, J. T. Nascent RNA Sequencing Reveals Widespread Pausing and Divergent Initiation at Human Promoters. Science 322, 1845–1848 (2008).

18. Seila, A. C. et al. Divergent transcription from active promoters. Science 322, 1849–1851 (2008).

19. Andersson, R. et al. An atlas of active enhancers across human cell types and tissues. Nature 507, 455–461 (2014).

20. Schmid, M. & Jensen, T. H. Transcription-associated quality control of mRNP. Biochimica et Biophysica Acta (BBA) - Gene Regulatory Mechanisms 1829, 158–168 (2013).

21. Takahashi, H., Lassmann, T., Murata, M. & Carninci, P. 5′ end– centered expression profiling using cap-analysis gene expression and next-generation sequencing. Nat Protoc 7, 542–561 (2012).

22. Thurman, R. E. et al. The accessible chromatin landscape of the human genome. Nature 489, 75–82 (2012).

23. Fejes-Toth, K. et al. Post-transcriptional processing generates a diversity of 5'-modified long and short RNAs. Nature 457, 1028–1032 (2009).

24. Wang, J. et al. Sequence features and chromatin structure around the genomic regions bound by 119 human transcription factors. Genome Research 22, 1798–1812 (2012).

25. Mathelier, A. et al. JASPAR 2014: an extensively expanded and updated open-access database of transcription factor binding profiles. Nucleic Acids Research 42, D142–D147 (2013).

26. Core, L. J. et al. Defining the status of RNA polymerase at promoters. Cell Reports 2, 1025–1035 (2012).

27. ENCODE Project Consortium. An integrated encyclopedia of DNA elements in the human genome. Nature 489, 57–74 (2012).

28. Trapnell, C. et al. Differential gene and transcript expression analysis of RNA-seq experiments with TopHat and Cufflinks. Nat Protoc 7, 562–578 (2012).

29. Andersen, P. R. et al. The human cap-binding complex is functionally connected to the nuclear RNA exosome. Nat Struct Mol Biol 20, 1367–1376 (2013).

30. Lin, M. F., Jungreis, I. & Kellis, M. PhyloCSF: a comparative genomics method to distinguish protein coding and non-coding regions. Bioinformatics 27, i275–i282 (2011).

31. Kaida, D. et al. U1 snRNP protects pre-mRNAs from premature cleavage and polyadenylation. Nature 468, 664–668 (2010).

32. Cooper, G. M. et al. Distribution and intensity of constraint in mammalian genomic sequence. Genome Research 15, 901–913 (2005).

33. Taylor, M. S. et al. Rapidly evolving human promoter regions. Nat Genet 40, 1262–1264 (2008).

34. Marques, A. C. et al. Chromatin signatures at transcriptional start sites separate two equally populated yet distinct classes of intergenic long noncoding RNAs. Genome Biol. 14, R131 (2013).

35. The FANTOM Consortium and the RIKEN PMI and CLST(DGT). A promoter level mammalian expression atlas. Nature 507, 462–470 (2014).

36. Kiss, T. & Filipowicz, W. Exonucleolytic processing of small nucleolar RNAs from pre-mRNA introns. Genes & Development 9, 1411–1424 (1995).

37. Koch, F. et al. Transcription initiation platforms and GTF recruitment at tissue-specific enhancers and promoters. Nat Struct Mol Biol 18, 956–963 (2011).

38. Kowalczyk, M. S. et al. Intragenic Enhancers Act as Alternative Promoters. Molecular Cell 45, 447–458 (2012).

39. Langmead, B., Trapnell, C., Pop, M. & Salzberg, S. L. Ultrafast and memory-efficient alignment of short DNA sequences to the human genome. Genome Biol. 10, R25 (2009).

40. Marstrand, T. T. et al. Asap: A Framework for Over-Representation Statistics for Transcription Factor Binding Sites. PLoS ONE 3, e1623 (2008).

41. Kim, H. & Park, H. Sparse non-negative matrix factorizations via alternating non-negativity-constrained least squares for microarray data analysis. Bioinformatics 23, 1495–1502 (2007).

42. Hoffman, M. M. et al. Unsupervised pattern discovery in human chromatin structure through genomic segmentation. Nature methods 9, 473– 476 (2012).

43. Ernst, J. & Kellis, M. ChromHMM: automating chromatin-state discovery and characterization. Nature methods 9, 215–216 (2012).

44. Quinlan, A. R. & Hall, I. M. BEDTools: a flexible suite of utilities for comparing genomic features. Bioinformatics 26, 841–842 (2010).

45. Dobin, A. et al. STAR: ultrafast universal RNA-seq aligner. Bioinformatics 29, 15–21 (2012).

46. Ramsköld, D., Wang, E. T., Burge, C. B. & Sandberg, R. An Abundance of Ubiquitously Expressed Genes Revealed by Tissue Transcriptome Sequence Data. PLoS Comput. Biol. 5, e1000598 (2009).

47. Cabili, M. N. et al. Integrative annotation of human large intergenic noncoding RNAs reveals global properties and specific subclasses. Genes & Development 25, 1915–1927 (2011).

48. Heinz, S. et al. Simple Combinations of Lineage-Determining Transcription Factors Prime cis-Regulatory Elements Required for Macrophage and B Cell Identities. Molecular Cell 38, 576–589 (2010).

